# Circulating extracellular vesicles from HIV-1 gp120-treated mice act as endogenous algogens, mediating and maintaining HIV-associated chronic pain

**DOI:** 10.1101/2025.02.07.637192

**Authors:** Subo Yuan, Jia Yi Liew, Jiani Bei, Ajay Pal, Yuan Qiu, Junkui Shang, Katherine Araya, Vivian Tat, Haiping Hao, Isaac Gamez, Connor Haines, Qing Chang, Tais B. Saito, Kamil Khanipov, Bin Gong

## Abstract

HIV-associated chronic pain (HIV-PAIN) remains prevalent in the post combined antiretroviral therapy era, affecting 30-60% of HIV patients worldwide. The underlying mechanism responsible for the development and maintenance of chronic pain remains unclear. gp120 is a causal factor of the HIV-PAIN and functions as an exogenous algogen. The pain experienced by human HIV-PAIN has been modeled in mice (referred to as mHIV-PAIN) using intrathecal (i.t.) injections of gp120. gp120 is a relatively short-term, static, exogenous algogen that is exhaustible *in vivo*. In authentic infection, HIV virions serve as the primary source of exogenous gp120, which initiates the early phase of clinical HIV-PAIN. Interestingly, while the source of replenishing gp120 decreases after antiretroviral therapy by suppressing viremia, the prevalence of chronic HIV-PAIN remains stable. To induce chronic pain in mice, gp120 needs to be repeatedly applied by the i.t. route. This raises a key question: Is an endogenous inexhaustible algogen responsible for maintaining the chronicity of HIV-PAIN?

In the present study, we isolated circulating small extracellular vesicles (sEV) from mice using our mHIV-PAIN model that is i.t. injected with gp120. We refer to such sEV as gp120-sEV herein. We observed that gp120 is absent in gp120-sEV. Following transfusion of gp120-sEV intrathecally, naïve recipient mice exhibit an extensive pain phenotype, including cold pain tested with we newly invented dry ice vapor cold test. RNA-sequence analysis suggests that gp120-sEVs induced expression of genes related to nociception and neuroinflammation pathways. These findings provide direct evidence that circulating sEV function as endogenous long-term “dynamic” algogens that enhance initial pain and extend the chronification of HIV-PAIN in mice, suggesting that chronic HIV-PAIN requires an exogenous algogen (gp120) paired with endogenous algogen (gp120-sEV), and that these components work synchronically to initiate and extend pain chronification. This double algogen concept provides a new insight into the pathogenesis of HIV-PAIN chronification. Our new mechanistic understanding will also assist in identifying new therapeutics to alleviate HIV-PAIN by targeting pathological gp120-sEV.

## Introduction

HIV-associated chronic pain (HIV-PAIN) remains prevalent in the post **C**ombined **A**ntiretroviral **T**herapy (cART) era, affecting 30-60% of HIV patients worldwide. This condition causes significant disability and reduced quality of life due to resistance to existing pain relief therapies compared with chronic pain in other diseases^1,2^. While cART effectively suppresses HIV viremia^3^, it does not eradicate the virus and the low-level viremia put one in the risk of pain chronification, whereby transient pain progresses into chronic, HIV-PAIN. the underlying mechanism responsible for the development and maintenance of chronic pain in remains unclear.

In chronic HIV-PAIN patients (referred as hHIV-PAIN), expression of HIV-1 gp120 is significantly higher in the spinal cord (SC) dorsal horn (SDH), the pain transmission center. This finding suggests that gp120 is a causal factor of the hHIV-PAIN and functions as an exogenous algogen^4^. The high level of gp120 could result from a latent viral reservoir in the central nervous system (CNS) that with the characteristic of the years-long (from 7 to 11.5 years) slow decay of HIV DNA (provirus) in the host genome. It forms the obstacle for HIV cure and provides the algogens for HIV-PAIN, despite the virus suppression achieved antiretroviral therapy (ART)^5-8^. The pain experienced by hHIV-PAIN has been modeled in mice (referred to as mHIV-PAIN) using intrathecal (i.t.) injections of gp120^4,9^. gp120 is a relatively short-term, static, exogenous algogen that is exhaustible *in vivo*. To induce chronic pain in mice, gp120 needs to be repeatedly applied by the i.t. route. In authentic infection, HIV virions serve as the primary source of exogenous gp120, which initiates the early phase of hHIV-PAIN. Interestingly, while the source of replenishing gp120 decreases after ART by suppressing viremia (to a nondetectable level of less than 1-5 copies/ml plasma)^6^, the prevalence of chronic hHIV-PAIN remains stable. This raises a key question: Is an endogenous inexhaustible algogen responsible for maintaining the chronicity of HIV-PAIN?

Extracellular vesicle (EV) are broadly classified into two categories, small EVs (sEV) (also known as exosomes, 50-200 nm) and large EVs^10,11^, distinguished by their particle sizes and biogenesis pathways^12^. The formation of sEVs is initiated by budding into the late endosome; the multivesicular body fuses with the cell membrane before releasing intraluminal vesicles as sEVs^12-14^. sEVs can be secreted from all cell types in the pain neurocircuit, including neurons^(7–9)^, astrocytes^(9–13)^, oligodendrocytes^(13–15)^, microglia^(16, 17)^, and Schwann cells^(18–20)^. sEV can also cross the blood-brain barrier (BBB) and blood-spinal cord barrier (BSB) to broadly and remotely target the pain neurocircuitry as mediators of intercellular communication^15-17^. sEV can convey signals to a large repertoire of neighboring and distant recipient cells by ferrying functional biomolecular cargoes, including proteins and nucleic acids, that contribute to diseases pathogenesis, and are being actively investigated^18^. In the context of neuropathic pain, most investigations have focused on comparing sEV cargoes, particularly with small noncoding RNA (sncRNA), and exploring their roles in *in vitro* models of neuroinflammation^19^. However, there is a notable lack of *in vivo* studies investigating the functional correlation between sEV and HIV-PAIN, as well as the identification of functional cargoes driving this process.

In the present study, we employed size exclusion chromatography (SEC) to isolate circulating sEV from mice using our mHIV-PAIN model that is i.t. injected with gp120. We refer to such sEV as gp120-sEV herein. We observed that gp120 is absent in circulating gp120-sEV. Following transfusion of gp120-sEV intrathecally, naïve recipient mice exhibit an extensive pain phenotype, and gp120-sEVs induced expression of genes related to nociception and neuroinflammation pathways. These findings provide direct evidence that circulating sEV function as endogenous long-term “dynamic” algogens that enhance initial pain and extend the chronification of HIV-PAIN.

## Results

### Quality assessment of circulating sEV from the plasma of gp120-treated mice showing chronic pain

Using SEC, we isolated circulating sEV from the plasma of mice on day 21 (D21) (mHIV-PAIN donor) post i.t. with recombinant gp120 (gp120-sEV) or heat-inactivated gp120 as control. Sizes and morphologies of isolated particles were initially evaluated using atomic force microscopy (AFM)^20-24^, zeta-potential measuring, and nanoparticle tracking analysis (NTA)^21,22^. We observed that sEV morphology (**Fig. 2A**) and size distribution (**Fig. 2B**) were not significantly altered between control- and mHIV-PAIN groups. Western immunoblot analysis showed that sEV expressed sEV markers, but not gp120 (**Fig. 3C**). The zeta potential, an indicator of surface charge and colloidal stability influenced by surface chemistry*^25^*, was also measured. We observed no significant differences between groups, suggesting that sEV from both groups are surface stable particles (**Fig. 3D**).

**Figure 1.**
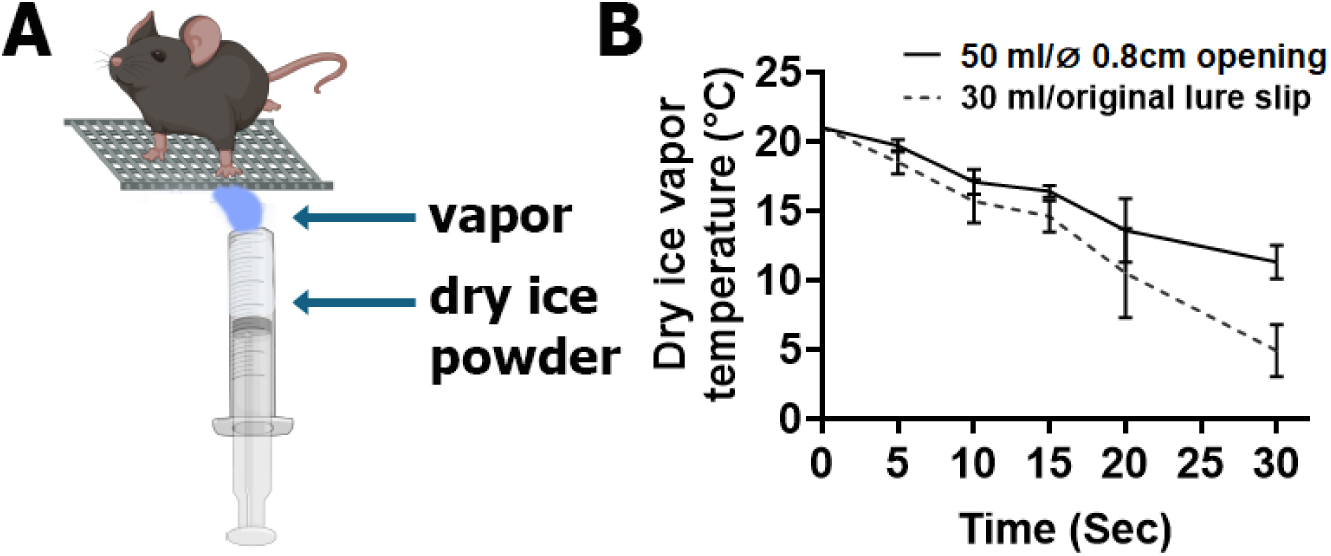
Dry ice vapor cold test measures hindpaw withdrawal latency as a measure of cold allodynia. A luer-lock syringe without the needle was cut at the tip and filled with a fine powder of dry ice, which could be extruded by pushing the syringe plunger. The evaporated dry ice vapor (blue) from the syringe immersed the mouse plantar and coolded it in from room T°21°C -17°C in 10sce, to 13.6°C in 20sec, to 11.3°C in 30sec. The distance between the syringe opening and the mouse hindpaw was 0.8 cm. Hindpaw withdraw latency was recorded as the time (in seconds) from application of the dry ice vapor until the hindpaw withdrawal happened.

**Figure 2.**
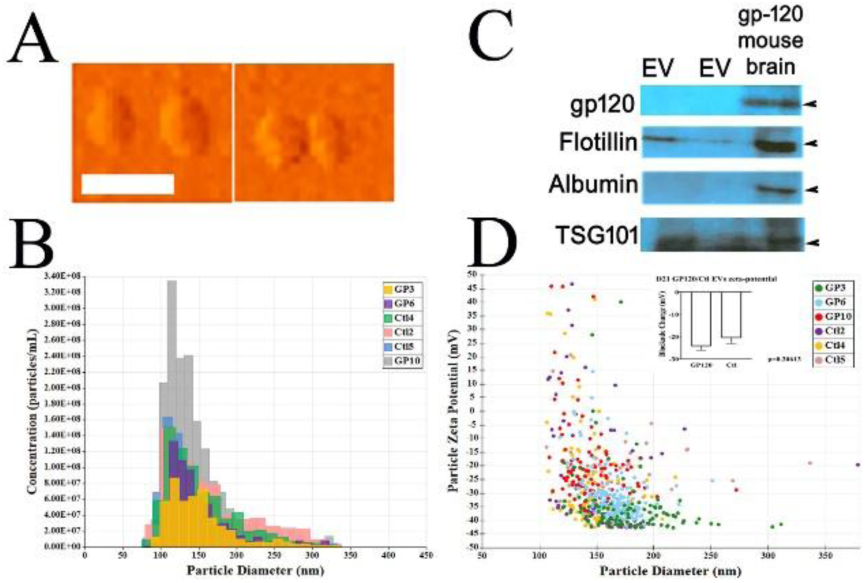
Characterization of sEV from the plasma of gp120-treated mice (gp120-EV) showing chronic pain (mHIV-PAIN) versus heat-inactivated gp120 mice (Ctrl-EV). **A**, Surface topographies of gp120-sEV and control sEV was verified using atomic force microscopy (AFM), scale bars, 200 nm. **B**, Vesicle size distribution of isolated EVs was analyzed using nanoparticle tracking analysis (NTA) (n=3 per group). **C**, Expression using western immunoblotting of indicated protein markers (gp120, flotillin, albumin, TSG101) in 100 μg protein of sEV or extracts from brain of a mHIV-PAIN mouse exhibiting chronic pain on Day 21. Samples in the two EV-lanes were control-sEVs (left) and gp120-sEVs (right), respectively. **D**, The zeta potential (in mV) was measured using the qNano-system, n=3 per group.

**Figure 3.**
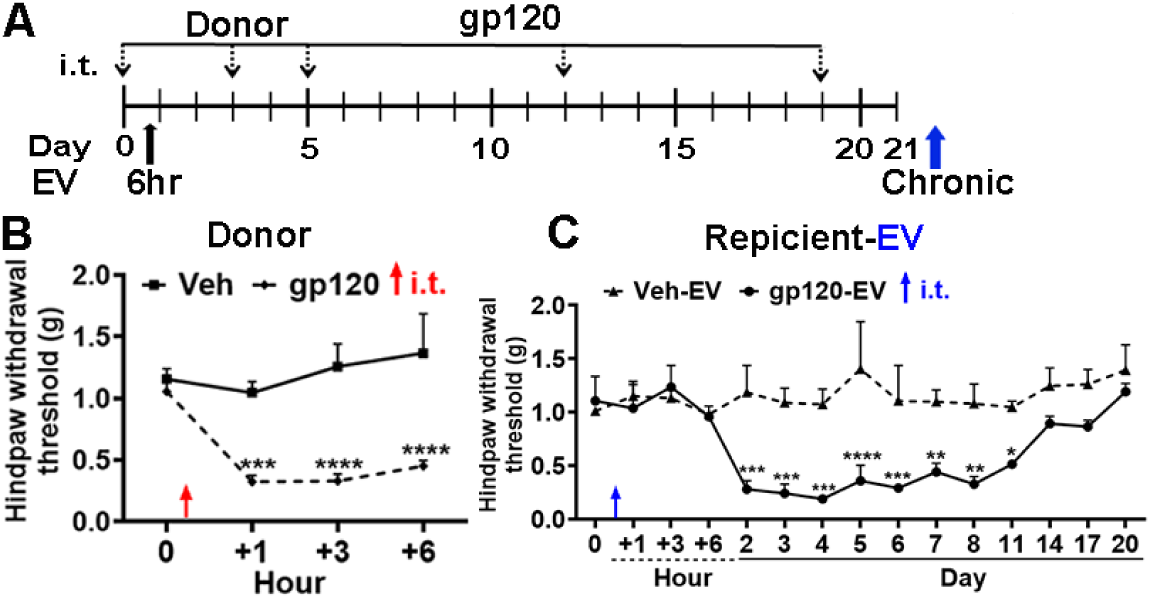
gp120-EV from acute mHIV-PAIN donor mice replicated pain in recipient mice. **A**, Dosing schedule of gp120 (i.t., 100 ng/20 g body weight [gbw]) in donor mice. **B**, Acute gp120 pain EV induced mechanical allodynia in recipient mice following one i.t. dose, 1.05×10^12^ particles/kg. Hindpaw withdraw threshold (PWT) (mean±SEM) was analyzed by two-way ANOVA with Bonferroni’s multiple comparisons test. *P<0.05, **P<0.01, ***P<0.001, ****P<0.0001. Donor mice: Veh (heat inactivated gp120), n=7; gp120, n=10. **C**, Recipient mice: Veh-EV, isolated from heat-inactivated gp120 treated donor mice, n= 6 and gp120-sEV, isolated from gp120 treated donor mice, n = 6.

### gp120-free, gp120-sEV works as an endogenous algogen to induce mechanical allodynia and temperature sensitivity in naïve recipient mice

gp120 is a well-reported algogen in both patients and mice^5,29^, but whether gp120-sEV is an algogen is unknown. One dose of gp120 (i.t., 100 ng/20 gbw) can induce acute allodynia lasting up to 5 days (**supplemental Fig. 1**) and repeated dosing of gp120 can induce “chronic” allodynia up to Day 21^5^. This work investigates the noxious effects of circulating gp120-sEV on recipient mice (**Fig. 3A**).

First, we tested whether gp120-sEV harvested from mHIV-PAIN mice 6 hr following a single dose of gp120 i.t. were able to induce nociception in naive recipient mice (**Fig. 3A**). We obtained 2.63×10^10^ particles of circulating sEV from an adult mouse (0.025 kg body weight, bw) on day 21 after i.t. gp120 administration. It should be noted that technically not all blood can be harvested by retro-orbital bleeding. Accordingly, a recipient mouse was transfused by the i.t. route with an sEV dose of 1.05×10^12^ particles/kg. We observed that recipient mice transfused with gp120-sEV did not show a nociceptive effect initially at 6 hr post i.t. dosing but began to present a typical allodynia on Day 2 post-i.t. injection. The allodynia lasted up to 11 days post-i.t. injection (Fig. 3B), supporting our hypothesis that gp120-sEV is a novel endogenous algogen.

We next sought to test the algogenic effect of gp20-sEV that was harvested on Day 21 post-dosing with five gp120 i.t. treatments (**Fig. 4A**). Distinct from the gp120-sEV harvested in the 6 hr group, gp120-sEV from mHIV-PAIN on Day 21 induced typical mechanical allodynia (**Fig. 4E**), as well as heat hyperalgesia and cold allodynia in donor mice (**Fig. 4F, G**). In recipient mice, the noxious effects associated with initiated severe allodynia started at 1hr post-i.t. dosing with gp120-sEV from Day 21 mHIV-PAIN mice, and last to 7 days without a trend of decreasing allodynia (**Fig. 4E**). gp120-sEV from the Day 21 group also induced heat hyperalgesia and cold allodynia in recipient mice (**Fig. 4F, G**), similar to donor mice treated i.t. with gp120 developed heat hyperalgesia (**Fig. 4C**). Regarding cold sensitivity test, hindpaw withdraw latency of donor mice increased, while in recipient mice it decreased, indicating that gp120 and gp120-sEV alternated the cold sensitivity differently. However, mice receiving a single dose of gp20-sEV (**Fig. 4E**) from Day 21 mHIV-PAIN donor mice that received five doses of gp120 by the i.t. route (**Fig. 4B**) demonstrated similar noxious effects, indicating gp120-sEV had stronger algogenic effect than gp120 itself.

**Figure 4.**
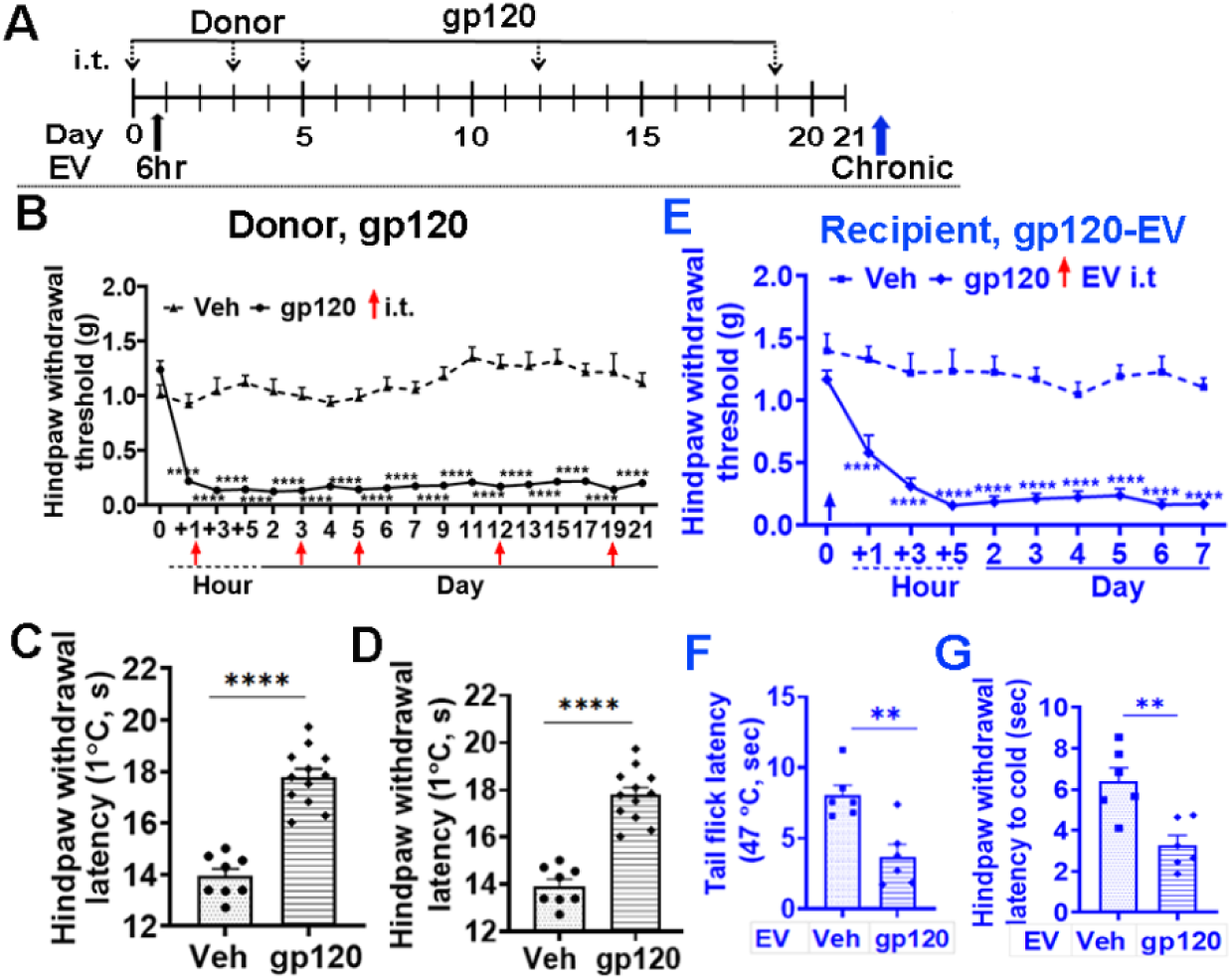
Chronic gp120-sEV replicated gp120 induced allodynia, cold allodynia and hyperalgesia in recipient mice. **A**, Dosing schedule of gp120 (i.t., 100 ng/20 gbw) in **mHIV-PAIN** donor mice **B**, Repeated i.t. injection of gp120 induced chronic mechanical allodynia in mice as measured by von Frey test and data expressed as PWT (in grams of force a of filament that elicits a withdrawal reflex). Veh, n= 11. gp120, n=6; two-way ANOVA test, ****P<0.0001. **C**, gp120 induced heat hyperalgesia. Mice number was same as A. Student t test. *P<0.05, ****P<0.0001. **D**, gp120 induced cold sensation alteration compared with Veh treated mice measured by hindpaw withdrawal latency elicited by dry ice vapor (hindpaw T°C drop range described in Fig. 1). Statistics same as B. **E.** gp120-sEV replicated mechanical allodynia in recipient mice; gp120-sEV was from **mHIV-PAIN** donor mice and Veh-sEV was from vehicle donor mice. Red arrow indicates time of administration of gp120-sEV or Veh-sEV, respectively. Veh, n=8; gp120-sEV, n=6. Statistical analysis was same as B. **F.** gp120-sEV induced heat hyperalgesia as measured by tail flick latency (in seconds at 47°C). Statistics is same as in C. **G.** gp120-sEV induced cold allodynia as measured by dry ice vapor test and hindpaw withdrawal latency was measured by seconds. Statistics is same as in C.

The comparative noxious effects between gp120 and gp120-sEV are summarized in **Table 1**. The table shows that a single dose of gp120 induces mechanical allodynia at 1hr post i.t. and last up to 5 days (**supplemental Fig. 1**) in donor mice. gp120-sEV collected at 6 hr from donor mice induced an initial hindpaw withdraw threshold (PWT) drop at 24 hr post i.t. in naïve recipient mice. This lower PWT (≤0.5 g) persisted up to 11 days. While 5 repeated doses of gp120 i.t. (**Fig. 3A**) maintained mechanical allodynia up to Day 21 in donor mice, and gp120-sEV from the Day 21 group caused an initial PWT drop at 1 hr post-i.t. in recipients, in which severe allodynia with low PWT ≤ 0.2 g persisted up to 7 days without decreasing. Collectively, these data suggest that 1.05×10^12^ particles/kg of gp120-sEV given by the i.t. route from the 6 hr group (single dosing in donors) has a longer algogenic effect in naïve recipient mice than does a single dose of gp120 in donor mice. While 1.05×10^12^ particles/kg of gp120-sEV from the Day 21 group has stronger algogenic effects in naïve recipients than multiple doses of gp120 in donor mice, this discrepancy indicates that gp120 and gp120-sEV induce pain via different nociceptive mechanisms.

### gp120-sEV algogenic effects depend on RNA cargoes

Exosomes can convey signals to a large repertoire of neighboring and distant recipient cells by ferrying functional cargoes. Although exosomes contain both proteins and RNAs, it has been documented that many effects of small EV on recipients are mediated by noncoding RNA cargoes^26-32^, particularly microRNAs (miRNA) and tRNA fragments^33-37^ at specific doses. RNAs were isolated from circulating gp120-sEV from mHIV-PAIN mice on Day 21. Using stem-loop RT-qPCR, which is a common method for detecting sncRNAs in EVs^33,38-40^, we measured exosomal expression levels of inflammatory relevant sncRNAs, including miR-23a, 24, 27, 30b, and tRFGly, which is reported to be enriched in circulating EV and associated with inflammation^24,41-43^, and miRNA let-7, a common microRNA in circulating exosomes^44,45^. We detected no differences in expression of these sncRNAs in sEVs (**Supplemental Fig. 2**).

Active permeabilization techniques have been widely employed in the field of sEV research, showing no significant impairment of sEV constitution, integrity, or functionality^48-50^. To determine the potential roles of RNA cargoes in gp120- sEVs, we employed electroporation-assisted active permeabilization^51,52^ by pretreating sEV cargoes with 20 µg/mL RNase. Such pretreatment completely erased the noxious effect of gp120-sEV (**Fig. 5**) in naïve recipient mice at 1.05×10^12^ particles/kg i.t, suggesting that the nociceptive cargo is RNA-cargo dependent.

**Figure 5.**
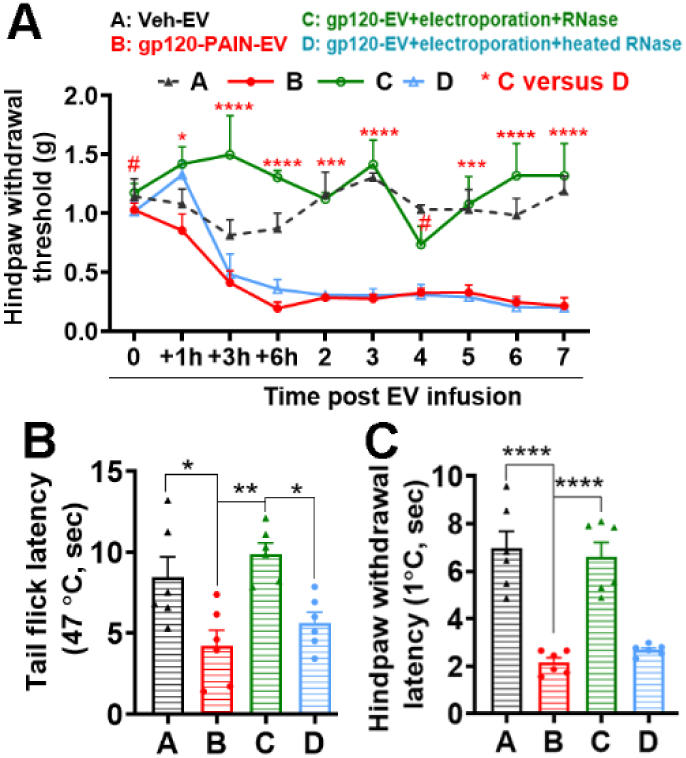
Noxious cargo of gp120-EV is RNA-dependent. **A**, gp120-sEV induced typical mechanical allodynia (red line) compared with Veh-EV(black dashed line). RNase digestion with 20 µg/mL gp120-sEV by electroporation resulted in a loss of the noxious effect (green line). The mock digested sEV retained its noxious effect to cause mechanical allodynia (blue line). **B**, RNase digested gp120-sEV lost its noxious effect to cause heat hyperalgesia, showing increased latency to heat stimulation in tail flick test (green column). **C**, RNAase digested gp120-EV lost its capacity to induce cold allodynia and restored hindpaw withdrawal latency to dry ice vapor in adult mice (green column). Each group, n=6. Data were presented as mean ± SEM, *p<0.05, **p<0.01, ***p<0.001, ****p<0.0001, 2-way ANOVA followed by Bonferroni’s multiple comparisons test in Graphpad Prism 8.0.2 A. one-way ANOVA followed by Holm-Sidak’s multiple comparisons test also was done using Graphpad Prism 8.0.2.

### gp120-sEV induced expression of nociceptive mRNA and relevant neuroinflammation pathways in gp120-sEV recipient mice

To evaluate neuroinflammatory responses, we created a heatmap to compare transcript expression levels of Ctrl-sEV (heat-inactivated gp120, 3 mice) and gp120-sEV-recipient mice (3 mice) (**Fig. 6A**). Overall, 19 genes were found to be upregulated in the spinal cords of gp120-sEV mice, while 13 genes were downregulated. We selected 36 pathways associated with neuroinflammation and nociception that were significantly different between the two groups for network analysis through Ingenuity Pathway Analysis (IPA) (**Fig. 6B**).

**Figure 6.**
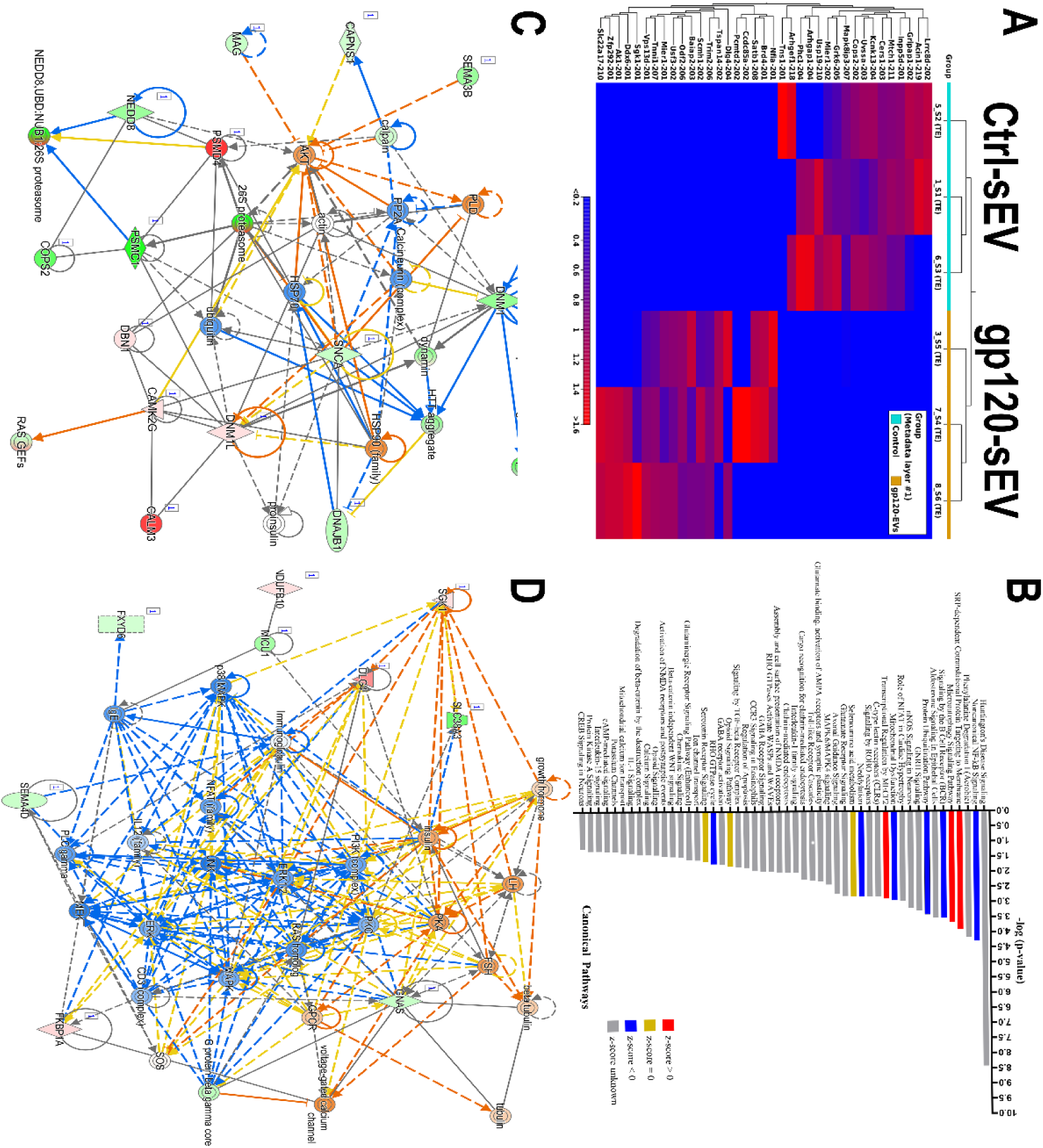
gp120-sEV recipient mice SC mRNA deep sequencing, transcript, and canonical pathway analysis. **A**, Heat map of transcript expression levels of SC from heat inactivated gp120-sEV (Ctrl-sEV, 3 mice) and gp120-sEV recipient mice (3 mice per group). **B**, Ingenuity Pathway Analysis (IPA) was used to analyze pathways associated with neuroinflammation and nociception in SC of gp120-sEV recipient mice. -Log value ≥2 was defined as significance alternation. **C** and **D,** The representative pathways identified through IPA from SC of gp120-sEV recipient mice that could be potentially regulated by miRNA expression in the SC of gp120-sEV recipient mice. Two networks were created **C**, Network 1: Downregulation (blue) of “calcineurin complex” and upregulation (yellow) of “AKT-actin-HSP90 pathway”. **D**, Network 2 was centralized with two opposite complexes and families: (1) Vertical upregulation pathways, “growth hormone-insulin” with three targets; “ERK1/2-JNK-PLC gamma” with two wing pathways, “RAS homolog” and “immunoglobulin”. (2) Horizonal upregulation pathways, “PKA-voltage-gated calcium channel”, and “PKA-insulin-SGK1”.

Fig. 6B shows that the Huntington’s Disease Signaling gene, HTT, was dramatically impacted in gp120-sEV recipient mice, with the -log p value up to 8.5. The HTT gene provides instructions for the huntingtin protein (HTT), which forms aggregates that disrupt neuronal function and eventually leads to death of neurons. HTT is a critical protein that senses and facilitates DNA damage repair during transcriptional elongation, which serves to improve genome integrity and neuronal survival^46^. However, we do not yet to know whether HTT is up- or down-regulated in gp120-sEV recipient mice. The dramatic gene alteration of HTT provides a unique opportunity for future studies, hinting at a novel mechanism relevant to neuroinflammation and pain.

Deep sequencing also revealed that the “SRP-dependent cotranslational protein targeting to membrane”, “microautophagy signaling pathway”, and “transcriptional regulation by MECP2” were significantly upregulated. It is worth noting that microautophagy paired with apoptosis is an adaptive response when neurons encounter stress or injury. Microautophagy activity assists with myelin clearance, promotes nerve regeneration, and attenuates pain, while attenuated apoptosis is responsible for the selective loss of GABAergic inhibition in SDH^47,48^. Upregulation of microautophagy may allow crosstalk with apoptosis in regulating proinflammatory cytokine expression in gp120-sEV mice. Increases in microautophagy were reported in DRG, spinal cord, and Schwann cells in mostly neuropathic pain rodent models, including chronic constriction injury, spinal nerve ligation, spared nerve injury pain rodent models^48^. Available data suggests that microautophagy represents an activated neuropathic pain and nerve injury cascade^49^.

The noncanonical NF-κB signaling pathway was found to be downregulated in gp120-sEV recipient mice. This pathway is upregulated in neuroinflammation and is linked with several chronic pain and nerve injury-induced pain rodent models^50^. The alteration of noncanonical NF-κB signaling in recipient mice did not align with its upregulation in donor mice and requires more study. Activation of “mitochondrial dysfunction” genes occur in neuroinflammation that affect cellular processes, such as calcium homeostasis and disrupted axon metabolism in donor mice^51^. It is possible that damage-associated molecular pattern was excluded in sEV cargoes in donor mice that led to mitochondrial dysfunction inhibition in recipient mice^52^.

The Rho family of GTPase genes (identified as “RHO GTPase cycle” gene on Fig. 6B was significantly inhibited in gp120-sEV mice compared to control-sEV mice. The Rho family of genes encodes proteins that belong to the Ras homolog family of guanosine triphosphate hydrolases that are important regulators in sensory neurons affecting neuronal axon guidance, cell migration and polarity, cell cycle progression, and regulation of gene transcription. These proteins robustly increase the transcriptional activity of NF-κB by phosphorylation of I-κB^53^. The mechanism correlated with inhibition of NF-κB signaling in recipient mice.

Other pathways related to immune responses, including “Toll-like Receptor Cascades”, “Interleukin-1 family signaling”, and “Chemokine Signaling”, were also identified by gene expression profiling in this study. In total, these data illustrate that circulating gp120-sEV can significantly affect signaling pathways relevant to neuroinflammation, immune responses, proinflammatory cytokine regulation, neuronal stress and injury, and sensory neuron axon guidance. We have clearly demonstrated that gp120-sEV is capable of transferring nociceptive and neuronal inflammation cargo molecules that contribute to the formation of neuroinflammatory-associated chronic pain.

To investigate the responses underlying the nociceptive effects of circulating gp120-sEVs, we next performed pathway analysis on SC tissues of gp120-sEVs mice harvested on day 21. A total of 480 pathways were identified using IPA of gene expression. We hypothesize that these signaling pathways are regulated by miRNA expression in gp120-sEV. Two networks were created. **Fig. 6C** illustrates Network 1, highlighting downregulation (blue lines) of the “calcineurin complex” and upregulation (orange lines) of the “AKT-actin-HSP90 pathway”. The calcineurin complex, along with PP2A, calpain, CAPNS1, HSP90, DNAJB1, HSP70, and ubiquitin, were identified as being inhibited in gp120-sEV mice compared to control mice. Calcineurin is a protein phosphatase whose primary target in T-cell activation is the dephosphorylation of nuclear factor of activated T-cells (NFAT), which regulates activation and control of thymocyte development, T-cell differentiation and self-tolerance^54,55^.

**Fig. 6D** illustrates Network 2, centralized with distinct complexes and families. The first pathway showing vertical upregulation (orange lines and circles) includes “growth hormone-insulin” with three targets (in blue): “ERK1/2-JNK-PLC gamma”, with two wings, “RAS homolog” and “immunoglobulin”. The second pathway (also in orange) is a horizonal upregulation pathway involving “PKA-voltage-gated calcium channel” and “PKA-insulin-SGK1” (in orange). Intraneuronal calcium increase is a critical mechanism for nociception coding^56^. Serum and glucocorticoid-inducible kinase 1 (SGK1) is widely expressed in the central nervous system and importantly regulates the sodium-chloride cotransporter and sodium-potassium gated channels ^57-59^. All these complex and targets activation are classic mechanisms in multiple pain models^60-62^, and were shown to be upregulated in gp120-sEV recipient mice in our study (**Fig. 6C**, orange color). Downregulated pathways include PI3K complex (in blue) with two related pathways: First, a vertical “ERK1/2-JNK-PLC gamma”, and second, a horizontal “p38 MAPK -NFAT (family)-JNK-MAPK-G protein beta gamma core”. Both the growth hormone-insulin and PI3K-ERK1/2 centralized networks are upregulated in different rodent pain models, while p-JNK, p-ERK and p-CaMKIIα were upregulated in mHIV-PAIN mice and HIV-infected patients with pain syndrome^4,63^ and downregulated in gp120-sEV-PAIN mice. We reason the nociceptive outcome in gp120-sEV mice is a result of the crosstalk between pathways and the interaction between up and downregulation of two major pain relevant pathways and networks. Notably, genes related to inflammatory signaling cascades, which contribute to up- and downregulation of processes in the pathogenesis of chronic HIV-PAIN, were intermingled in the spinal cords of gp120-sEV mice compared to control-sEV mice. However, this outcome solely resulted from the gp120-sEV cargoes that transferred nociceptive cargoes from donor to recipient mice, resulting in the nociceptive phenotype in naïve recipient mice.

Combining the analyses of transcriptomic, mRNA-seq, and canonical pathway data, it is evident that gp120-sEV cargoes act as secondary algogens that are generated from the primary gp120 algogenic effect and have the characteristics of endogenous algogens capable of transporting pro-inflammatory RNA cargoes. The cargoes modulate neuroinflammatory signaling pathways within pain circuits and drive the chronification of HIV-PAIN. In summary, this study provides evidence that the primary gp120 viral coat protein initiates pain and acts as an exogenous algogen, while the EV cargo of gp120 (i.e., gp120-sEV) can initiate and extend pain, functioning as an endogenous algogen leading to chronic HIV-PAIN. Furthermore, we postulate that the two types of algogens need to work synchronously.

Collectively, the bioinformatic findings presented provide mechanistic insights into the role of gp120-sEVs in HIV-PAIN pathogenesis and highlight specific pathways and molecular targets that may inform future studies. In addition, gp120-sEV may serve as a potential target of therapeutic intervention.

## Discussion

In this work, we reported for the first time a critical algogenic role for gp120-sEV derived from HIV-PAIN mice that function as endogenous algogens to induce pain, and, most importantly, to extend pain chronification in recipient mice. HIV-1 gp120, derived from HIV infection, works as an exogenous algogen to initiate HIV-PAIN in donor mice. In our model, gp120-sEV recipient mice resemble gp120 donor mice in that they fully exhibit mechanical allodynia (as measured by the Von Frey test and PWT), heat hyperalgesia (as measured by tail flick latency), and cold allodynia (as measured by the dry ice vapor cold test). Unexpectedly, gp120-sEVs possessed stronger nociceptive efficacy than gp120 itself. Our HIV-PAIN mouse model demonstrated that exogenous gp120 algogens initiated pain and shed gp120-sEV, while the endogenous gp120-sEV extended pain and contributed to chronic pain.

This study provides a unique insight into the mechanism driving the high prevalence of HIV-PAIN worldwide despite HIV viral suppression by ART^2,3^. Spinal cord expressed gp120 is a well-recognized algogen in rodents and humans. Under viral suppression, a slow decay of residual viremia and latent reservoirs in the CNS (due to suboptimal drug penetration) constitute the source for replenishing gp120, which is limited due to successful ART^6,7^. The existing gap in our knowledge is how gp120 continuously exerts its nociceptive potential under the stress of ART. This study demonstrated that circulating gp120-sEV from HIV-PAIN donor mice was gp120-free, but ferried enough noxious cargo to induce the full pain phenotype in naïve recipient mice. It is worth noting that gp120-sEV can wisely avoid being targeted by ART, but possesses a stronger algogenic capacity to act as an ART-exempted, endogenous algogen (**Fig. 7**). Gp120-sEV initiates pain in naïve mice, enhances pain in feedback way and extends pain chronification in mice. This study provides novel insights into why HIV viremia can be well-controlled by ART, while HIV-PAIN cannot, and chronic pain persists with relatively high prevalence in those individuals on ART therapy.

**Fig 7.**
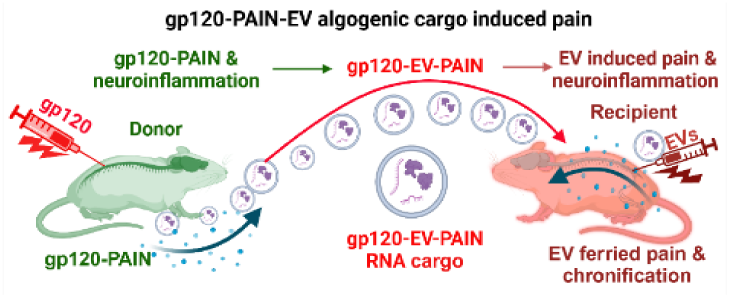
Schematic diagram depicting gp120-sEV as endogenous algogens, ferrying algogenic cargo, and replicating pain in naïve recipient mice. HIV-1 gp120 caused pain in donor mice as a known algogen. The donor mice derive gp120-sEV, which ferries nociceptive cargo to the pain neurocircuitry of naïve recipient mice to cause pain. simultaneously, gp120-sEV exerts nociceptive effects in a feedback loop manner, enhancing pain and maintaining pain chronification in donor mice. Therefore, gp120-sEV is a novel endogenous algogen.

To fully describe gp12-sEV induced pain in our mouse model, we invented a novel “dry ice vapor cold test” and employed it to measure cold allodynia. This method, used in pain behavior testing, is first reported here. The existing cold tests seek to apply solid cold material (e.g., acetone drop, cold plate or glass) directly to rodent hindpaw plantar skin and record the withdraw latency in seconds, also observing noxious behavior signs such as shake, filch, brisk, jump, or from cold stimuli^64-68^. The disadvantage of this approach is that mouse skin touching an extremely low temperature material triggers an immediate, conditional withdraw reflex, instead from authentic transduction of cold nociceptors. The additional disadvantage is that the touch and withdraw conditioned reflex in the cold test makes mice sensitive to repeated measurements due to memory and quick learning^69^. Although efforts were made to reduce this disadvantage, the distinct disadvantage still limit its use^70,71^. The advantage of the “dry ice vapor cold test” is the use of cold vapor that cozily and softly touches the plantar of the hindpaw. The vapor test avoids the solid cold material touch and withdraw reflex, makes better repeated tests possible, and results in stable withdraw latency between experiments. This novel technique adds a new method to cold pain test in basic pain research and is especially helpful in existing rodent pain model that more sensitive to cold, spinal cord and nerve injury, chemotherapy-induced pain and neuropathy, joint arthritis induced pain, and type of neuropathies.

Since this work focuses on the chronicity of HIV-PAIN, we first need to define what is the chronicity of HIV-PAIN. The clinical definition for chronic pain is pain lasting more than 3 months^72-74^. However, basic studies of pain have no such definitive definition. In this study, we define acute and chronic pain depending on which nociceptor mediates pain. gp120 spinal exposure leads to nociceptor cutaneous innervation inflammation and degeneration in mice hindpaw plantar skin, an area where von Frey stimulates applied on to obtain hindpaw withdraw threshold (PWT). The innervation of protein gene product 9.5 (PGP9.5^+^) is measured by intraepidermal nerve fiber (IENF). gp120 treatment for 1 to 14 days causes PGP9.5^+^nociceptor inflammation; at this stage, the pain mediated by the PGP9.5^+^ nociceptor is acute pain. Day 14 is the transitional point between acute and chronic pain. After 14 days, the predominate PGP9.5^+^ nociceptor is close to gone, and pain enters the chronic stage^4^. For this reason, in this study acute gp120-sEVs were collected from HIV-PAIN mice at 6hr following gp120 treatment, while chronic gp120-sEV were collected on day 21.

Chronic HIV-PAIN requires two types of algogens, exogenous gp120 for initiation and endogenous HIV-sEV for pain chronification. gp120 expressed in patient SDH forms the “first hit” in the pain circuit^4^. Afterwards, gp120-sEV are shed into the blood stream from gp120 treatment, where they function as endogenous algogens to form “secondary hits” on the pain neurocircuit in a positive feedback manner, independently initiating, maintaining, and extending pain chronification as a full algogen (**Fig. 3C, 4E**). To elucidate the nociceptive roles of the two types algogens, two different pain mouse models are needed. The advantage of our mHIV-PAIN mouse model is that it allows investigation of the algogenic role of gp120 without interference by other HIV-viral components^4^. The advantage of our HIV-sEV mouse model is that it allows us to focus on the noxious effect of HIV-sEV. The creative pairing of HIV-PAIN and HIV-sEV pain mouse models enables us to differentiate the noxious role of the exogenous algogen in gp120 donor mice and the endogenous algogen in gp120-sEV recipient mice. The existing gp120 pain rodent models developed by spinal cord, DRG, and sciatic nerve gp120 exposure are used to mimic HIV-PAIN and investigate neuroinflammatory mechanisms and various interventions^63,75-77^. Importantly, we reported that the mHIV-PAIN mimics the pain and spinal cord molecular changes of hHIV-PAIN patients and confirmed that gp120 is an exogenous algogen in both rodents and humans^4,78^. Humanized mice possess a functional human immune system and accurately recapitulate HIV infection, making it a preferred HIV infection rodent animal model. Incidentally, ecoHIV mice also develop latent viral reservoirs and are a well-accepted rodent HIV infection model^79-81^. Additional consideration is sEV shed from HIV infected cells have the same size, biogenesis, and physical properties as HIV virus and are indistinguishable each other. HIV-shed sEV contain host viral genetic RNA and proteins other than gp120^82,83^. This characteristic limit the application of HIV infection of humanize mice in HIV-sEV pain studies. As we focus only on the noxious effect of pathogen-free sEV and do not expect sEV mixed with HIV virions or sEV to contain HIV viral components. We chose the mHIV-PAIN pain mouse model to generate gp120-sEV that we show to be free of gp120 (**Fig. 2C**), which are perfect for our nociception study. Previous work intermingled the noxious effects of gp120-sEV with gp120 in a single pain mouse model, which impeded the discovery that gp120-sEV is new type of algogen. We reported on the pathological effect of circulating sEV isolated from Alzheimer’s patients on recipient endothelial cells and found that the Alzheimer patient sEV weakened the integrity of the blood-brain barrier in a recipient three-dimensional microvascular tubule formation model^84^. Testing the pathological role of sEV from donors in recipient mice or cells is a creative approach that allows us to untangle intermingled mechanisms occurring in the body. In this study, our models were employed to differentiate the noxious role of exogenous gp120 and endogenous gp120-sEV as algogens.

In this study, we found that gp120-sEV showed higher algogenic efficacy than gp120 itself. To maintain a chronic pain up to day 21 requires that gp120 be injected five times by the i.t. route (**Fig. 4A, B**). Acute pain sEV were harvested at day 6hr following a one-time i.t. injection of gp120. Acute gp120-sEV at a dose of 1.05×10^12^/kgbw (**Fig. 3B**) that is approximately equal to sEV in circulating mouse blood^85^. This dose of sEV maintained high level chronic pain with low intensity of threshold till to day 11 after acute gp120-sEV (**Fig. 3C**).

We reported that HIV-PAIN patients on ART exhibit a 10 times higher level of gp120 in the SDH compare with HIV patients without pain, and the expression was irrelevant to HIV plasma virion load^4^. HIV RNA in plasma ranging from 1 to 1000 copies/ml plasma depend on individual regimens and correlate to patients’ pre-ART viremia loads. The half-life of viremia is 3.7 to 11.7 years, and viremia results from the dynamic of HIV DNA provirus and long-lived viral reservoirs resulting from suboptimal drug penetration, especially in the CNS^5,7^. These reports suggest that persistent ART regimens can gradually decrease viremia even though SDH algogenic gp120 was kept at high levels, resulting in active shedding of gp120-sEV.

To compare the algogenic effects of gp120 with gp120-sEV, a key factor is dose. The sEV dose given to recipient mice was calculated based on the following considerations. The blood volume of an adult mouse (bw 0.025kg) is approximately 1.46 ml, and the blood volume we collect is 700-1000 µl max. Fifty-four percent of plasma can be recovered from whole blood, thus the sEV particles obtained from plasma ranged from 9.6×10^11^ to 6.5×10^12^ /kgbw^85^. The key goal of the present study is to investigate whether gp120-sEV is an endogenous algogen. Therefore, based on the range of circulating sEV particles, we used the median sEV particle number of 1.05 ×10^12^/kgbw for use in a single i.t. injection. This dose is approximately equal to the EV concentration circulating in a mouse. One limitation of this study is that the actual number of circulating sEV reaching the pain circuit remains unknown but will be investigated in future studies. We also plan to perform *in vivo* sEV dose-dependent trials involving different delivery methods to address the algogenic efficacy of gp120 and gp120-sEV.

To fully investigate gp120-sEV-induced molecular changes involving neuroinflammation and nociception in recipient mice, we utilized mRNA bioinformatics to provide insights on genes that are be negatively downregulated by miRNA cargoes of gp120-sEV. Neuroinflammation, immune responses, neurodegenerative processes, voltage gated potassium channel, neuronal sensitization, and nociception transmission are pertinent to chronic pain, and were found to be altered. The most interesting finding identified in Network 1 (**Fig. 6C**) is the inhibition of a pathway pertinent to chronic pain (the calcineurin complex along with the PP2A-calpain-CAPNS1-SHP90 [family]-DNAJB1 pathway). This pathway is also involved in HSP70 and ubiquitin, which together inhibit HTT aggregates. The central complex includes calcineurin, and immunosuppressing it with calcineurin inhibitors such as cyclosporine and tacrolimus in organ transplant patients caused calcineurin inhibitor pain syndrome that involves severe bilateral hip and lower extremity pain^86-89^. Extrapolation of these effects indicates that downregulation of calcineurin may exert a nociceptive effect in our gp120-sEV mouse model. The calcineurin complex also indirectly activates the protein kinase B(AKT) group, which are critical protein kinases in cell survival and growth in SDH; a pain transmission center; and development and maintenance of several chronic rodent pain models, such as diabetic neuropathy, neuropathic pain, bone cancer pain, capsaicin-induced pain, and opioid-induced hyperalgesia^60,89-92^. In our pathway, AKT connects with PLD-HSP90 and HSP70. The inhibition of calcineurin and activation of AKT provides *in vivo* evidence that these gene products play a crucial role in gp120-sEV-induced neuroinflammation and nociception.

Another interesting finding is the highly significant (-log p value close to 8.5) alternation of HTT signaling, hinting that HTT signaling may be a novel mechanism relevant to pain and neuroinflammation. To our knowledge, in addition to inherited gene mutation (CAG trinucleotide repeat) caused HTT variant that caused death of neurons in stratal neurons and induced motor, cognitive and emotional-volitional personality sphere disorders ^93^. HTT was also found to be a critical protein that senses and facilitates DNA damage repair during transcriptional elongation, which improves genome integrity and neuron survival^46,94^. To our knowledge, there are no reports that HTT signaling is involved in nociception-related neuroinflammation. Thus, this alteration may indicate that DNA integrity that is long been ignored in pain research, could be a novel mechanism for HIV-PAIN.

We observed two unexpected alternations in the spinal cord of gp120-sEV mice. In one pathway analysis, “noncanonical NF-κB signaling” was found to be downregulated in the spinal cord of gp120-sEV recipient mice (**Fig. 6B**, blue color). This pathway is upregulated in neuroinflammation and linked with several chronic and nerve injury pain rodent models^50^. However, this alteration did not align with upregulation reported in pain mouse models. In Network 2, two downregulated pathways were observed: (1) vertical pathway PI3K-ERK1/2-JNK-PLC gamma and (2) horizontal pathway centralized with p38MAPK, IgE, PLC gamma, MEK, CD3 complex, NFAT (family), MAPK-G protein beta gamma (**Fig. 6C**, blue color). PI3K centralized networks were upregulated in different rodent pain models and p-JNK, p-ERK and p-CaMKIIα were upregulated in a gp120 acute pain mouse model and HIV-infected patients with pain syndrome^4,63^. Li *et al.* reported that 12 hr gp120 treatment activated the Wnt5a/CaMKII and Wnt5a/JNK signaling pathways in the SC ^63^. We reported that p-JNK inhibitor sp600125 blocked gp120-induced allodynia in mice up to 6 days^78^. The anti-PI3k/Akt pathway is a potential chronic pain therapeutic target^61^.

Pathological sEV induced mRNA downregulations is also reported in other rodent pain models. Luo *et al.*reported the downregulated expression of immune response gene in recipient microglia uptaken sEV from spare nerve injury (SNI) pain mouse, upregulated gene expression in microglia uptaken sEV from sham mice, Luo et al suggest SNI-induced nerve injury may contribute to the inhibition^95^ In our study, miRNA present in RNA cargoes may negatively regulate mRNA expression in the spinal cord of recipient mice. Although the noncanonical NF-κB signaling and PI3K complex were downregulated in network 2, however, the growth hormone-insulin centralized pathways are upregulated in the same network, which positively contribute to neuroinflammation and nociception. The overall effect of upregulation and downregulation causes neuroinflammation and induces chronic gp120-sEV-assiciated pain.

The present study verified that the noxious effect of gp120-sEV is from RNA cargoes. We have not yet sequenced the miRNA or analyzed targets for experimental validation. Nevertheless, this limitation does not affect the concept that gp120-sEV is a novel endogenous algogen that is able to initiate, enhance, and extend pain chronification. Our future work will continue to utilize the gp120-sEV pain mouse model with sEV dose response, targets validation and miRNA sequence.

Based on the outcome of the study, we propose that chronic HIV-PAIN requires an exogenous algogen (gp120) paired with endogenous algogen (gp120-sEV), and that these components work synchronically to initiate and extend pain chronification. This double algogen concept provides a new insight into the pathogenesis of HIV-PAIN chronification. Our new mechanistic understanding will also assist in identifying new therapeutics to alleviate HIV-PAIN by targeting pathological gp120-sEV.

## Materials and Methods

Adult C57BL/6J mice, 2-4 months old with body weight 18-22 g, were purchased from The Jackson Laboratory, cat# 000664. Mice were housed in cages with soft bedding under a 12-hour reverse light/dark cycle. Animal procedures were performed following protocols that were reviewed and approved by the University of Texas Medical Branch Animal Care and Use Committee. Every attempt was made to reduce the number of animals used in these studies.

### Reagents

HIV-1 gp120 protein (HIV-1 gp120Bal, Cat # ARP-4961 produced in HEK293 cells), batch number 160227 and ARP-11437 anti-HIV-1 gp120 monoclonal (39F) antibody were obtained through the NIH AIDS Research and Reference Reagent Program, Division of AIDS, National Institute of Allergy and Infectious Diseases, NIH. Unless otherwise indicated, all other reagents were purchased from Thermo Fisher Scientific.

### Gp120 i.t. injections

Intrathecal injections were used to deliver gp120 protein or gp120-sEV to mice as we described^4,78^. Mice were anaesthetized by placing them inside a chamber connected to valproate isoflurane 3%-2.5% mixed oxygen with at flow rate of 1 L/min. A 30.5-gauge stainless steel needle attached to a 10 µl luer-lock tip syringe (Hamilton, Reno, NV) was used to perform the i.t. injection. To insert the needle into the spinal cord, the intervertebral space between L5 and L6 was located with one hand, and the needle was inserted into the intervertebral space at a 45° angle and injected with the other hand. A successful intrathecal placement of the needle tip was judged by a tail twitch^4,78,96^. gp120 protein was dissolved with 0.1 % bovine serum albumin (BSA) in 0.1 M phosphate-buffered saline (PBS) at a final concentration of 15 ng/µl, and the injection volume was calculated based on the body weight of the mouse.

### Von Frey mechanical sensitivity test

The Von Frey test was used to measure mechanical allodynia and performed as we described previously ^64,78,97^. Briefly, mice were habituated within a Van Frey plexiglass box with a mesh bottom in the morning for three continuous days prior to the behavioral test. On the test day, mice were restricted in the box for 20 min to acclimatize them to the test condition. The test was performed by using filaments of varying gauges (Stoelting, Wood Dale, IL) to perpendicularly, gently, and stably point to the center of hindpaw plantar mouse skin with appropriate force until the filament was slightly bent at the touch end and remained stable under force for about one second. The force exerted from the von Frey filament was calibrated depending on their stiffness. Stimulation began with the filament of 3.61 (0.4g) followed with up (increased) and down (decreased) stimulation force to obtain three effective pairs of stimulation response^98-100^. Signs of noxious withdraw response included abrupt perpendicularly and horizontal withdrawal of the hindpaw. The threshold of the force (in g) that evoked a noxious response was computed as mean ± SE and plotted with GraphPad Prism 8. Statistical significance was determined using one-way or two-way analysis of variance (ANOVA) with post-hoc tests.

### Tail flick test

The tail flick test is used to assess heat hyperalgesia in mice by measuring the latency period after applying heat stimulation. This is done by immersing a mouse’s tail in a predetermined temperature^64,101^. The mouse is gently and securely restrained in a triangle-shaped plastic bag, with the acute angle end open to allow the tail to extended outside^102^. This setup allows the experimenter to safety hold the bag and immerse the mouse tail in a warm water bath with a precise temperature setting. The tail is submerged both to provide a heat stimulus and to record the time it takes for the mouse to flick its tail away from the heat. The distal half of the mouse’s tail is immersed in a water bath (46–52°C), optimizing the temperature to achieve accurate withdrawal latency without causing agitation or inaccurate results. The latency period (in seconds) is recorded from immersion to the obvious tail flick. Data was expressed as mean ± SE and statistical significance was determined by student t test.

### Dry ice vapor cold test

Cold allodynia is one of the characteristics of inflammatory and neuropathic chronic pain that the type of HIV-PAIN belongs^103^. The first author developed the “dry ice vapor cold test,” as illustrated in **Fig. 1**. Initially, mice are placed inside a transparent, ventilated Von Frey test chamber with a mesh bottom for one hour each day over three consecutive days to acclimatize the mice to the experimental setup. Next, a barrel from a 50 ml or 30 ml syringe is filled with crushed dry ice powder, and the plunger is pushed to compress the powder to the tip end. The luer lock tip of the syringe was cut to create an opening for dry ice vapor release. This 50 ml syringe, with a 0.8 cm diameter (⌀) opening, is optimal for releasing a concentrated vapor flow that softly touches the hindpaw, avoiding a too concentrated hard vapor flow point released from the build-in syring tip that could induce a conditional reflex. The experimenter holds the syringe under the mesh, maintaining a distance of 0.8 cm from the plantar hindpaw, ensuring the vapor fully contacts the plantar glabrous skin without physical touch. Thus, the cold vapor serves as a direct cold stimulus to the hindpaw. The syringe is moved away immediately upon hindpaw withdrawal, and the test is terminated if no response is observed within 30 seconds to prevent cold injury. The latency of hindpaw withdrawal is recorded in seconds using a stopwatch. The latency values were computed as mean ± SE and plotted using GraphPad Prism 8. Statistical significance was determined using a student’s t test.

We optimized two sizes syringes (30ml and 50ml) and observed the temperature(T°) drop range. The 50ml syringe 0.8cm diameter opening in the lure slip area released vapor T° cooled the bulb of thermometer from room T° 21°C to 17°C in 10sec, to13.6°C in 20sec, to 11.3°C in 30s. 2). The 30ml syringe with original lure slip released vapor temperature cooled the bulb of thermometer from 21°C to 15.7°C in 10sec, to 10.5°C in 20sec, to 4.9°C in 30s. The 30ml syringe dropped T° quicker than the 50ml syringe, but it caused mice conditional hindpaw withdraw reflex due to stronger focused cold vapor. As most mice response to the vapor cold stimuli within 20sec, therefore, the vapor T° of 50ml syringe dropped from 21°C to 13.6°C in 20sec, that fallen in definition when cold T° lower than 15°C, should be count t as cold allodynia test.

### Mouse blood collection

Mice were terminal anesthesia 3% isoflurane in oxygen) and restrained by hand, with the mouse’s neck gently scruffed so that the eye was made to bulge. The eye was removed using sharp tipped forceps and blood was collected in a 1.5ml Eppendorf tube coated with 0.5M ethylenediaminetetraacetic acid (EDTA; pH 7.4; cat no. 0215252290, MP Biomedicals. Eppendorf tubes were coated with EDTA solution upon repeated dispensing and aspirating of 1ml of 0.5 M EDTA solution under sterile conditions five to six times, after finishing coat, left 10µl EDTA solution inside tube.

### SEC and NTA

sEV particles were isolated using size exclusion chromatography. Filtered mouse plasma was placed onto a qEVoriginal column (Izon, New Zealand). Fractions 7 to 9 were collected as the sEV-enriched fractions and concentrated using 100,000 MWCO PES Vivaspin centrifugal filters (Thermo Fisher Scientific)^21,24^. To treat EV-cargos with RNase, sEV samples (4 x 10^7^ particles/mL) and RNase (20 µg/mL) (Thermo Fisher Scientific) were mixed in 50 mM trehalose (Thermo Fisher Scientific) pulse buffer according to the manufacturer’s instructions. Suspended sEV particles were electroporated in 4 mm path length electroporation cuvettes using a BTX ECM 399 system (Harvard Apparatus, MA) with a voltage of 400V and a time constant as 5 ms. sEV samples were concentrated using 100,000 MWCO PES Vivaspin centrifugal filters prior to downstream applications.Nanoparticle tracking analysis (NTA) was performed using the tunable resistive pulse sensing (TRPS) technique and analyzed on a qNano Gold system (Izon, Medford, MA) to determine the size, concentration, and zeta-potential values of sEV particles. CPC200 calibration particles (Izon) were diluted in filtered Izon-supplied electrolyte at 1:100 to equilibrate the system prior to measuring using the manufacturer’s instructions. Measurements were conducted using NP100 nanopores (analysis range: 50–330 nm, Izon). A stable current of approximately 140 nA was achieved by adjusting the voltage. To determine the zeta potential of sEVs, nanopores were calibrated at three distinct voltages (V1 = 0.88 V, V2 = 0.7 V, V3 = 0.5 V) and four pressure settings (P0 = 0 mbar, P1 = 1 mbar, P2=5 mbar, and P3=10 mbar). Zeta-potentials of samples were measured at V1 and P0. For size distribution measuring, P2 and P3 were applied. The size, density, zeta-potential, and distribution of the particles were analyzed using qNano software (Izon). Imaging of label-free extracellular vesicles using AFM as we described^21,23^.

### Western immunoblotting

Western immunoblotting was performed as previously described^4^. Briefly, harvested sEVs were mixed with radioimmunoprecipitation assay lysis buffer (1% Nonidet P-40, 50 mM Tris-HCl, 0.25% Na-deoxycholate, 150 mM NaCl, 1 mM ethylenediaminetetraacetic acid, pH 7.4) containing a protease inhibitor cocktail (Sigma, St Louis, MO). After centrifugation (12,000 x *g*, 10 minutes), the supernatant was collected, and the protein concentration was determined using the BCA Protein Assay Kit (Pierce Biotechnology, Rockford, IL). Equal amounts of protein were separated by electrophoresis using sodium dodecyl sulfate–polyacrylamide gels, followed by transfer to polyvinylidene fluoride membranes. Following protein transfer, the membranes were washed once in Tris-buffered saline containing (TBST; 50 mM Tris, 150 mM NaCl, 0.05% Tween 20, pH 7.6) and then incubated in blocking buffer (TBST buffer with 5% nonfat milk powder). The membranes were then incubated sequentially with primary and secondary antibody. Protein bands were visualized using the Enhanced Chemiluminescence kit (Pierce) and quantified using NIH ImageJ software.

### Statistical analysis

All data were presented as mean± standard error of the mean and analyzed using SPSS version 22.0 (IBM^®^ SPSS Statistics, Armonk, NY). A two-tailed Student’s *t*-test or one-way analysis of variance (ANOVA) was used to explore statistical differences. If the ANOVA revealed a significant difference, a post-hoc Tukey’s test was further adopted to assess the pairwise comparison between groups. The level of statistical significance for all analyses was set at *p*<0.05.

### RNA sequencing and bioinformatic analysis

Total RNA was prepared from spinal cord tissue samples of control-sEV- and gp120-sEV recipient mice using TRIzol™ (Thermo Fisher Scientific), and total RNA was quantified using a NANODrop™ 2000 spectrophotometer.

The mRNA sequencing library was prepared using NEBNext® Poly(A) mRNA Magnetic Isolation Modules and NEBNext® Ultra™ II Directional RNA Library for Illumina preparation kits, according to the manufacturer’s protocol. The libraries were pooled and sequenced using an Illumina NextSeq 2000 instrument with a P1 100 cycle flowcell for single end 100 bp reads. RNA-Seq was performed by the Next Generation Sequencing Core at the University of Texas Medical Branch at Galveston.

CLC Genomics Workbench v24.0.2 was used for bioinformatical quality control and mapping of the RNA-Seq data. Sequencing data was initially trimmed using CLC’s Trim Reads 2.9. Reads containing nucleotides below the quality threshold of 0.05 (using the modified Richard Mott algorithm), those with two or more unknown nucleotides, or sequencing adapters were left out of the analysis. Filtered sequencing reads were then processed using the RNA-Seq Analysis 2.7. Reads were mapped using a local alignment strategy against the mouse GRCm39 reference genome with annotation version GRCm39.111 to identify gene and transcript abundance. Differential gene expression analysis was performed as part of the Differential Expression for RNA-Seq v2.8. The gene and transcript expression count were modeled by a separate Generalized Linear Model (GLM), and statistical significance was evaluated using the Wald Test. Significance was determined using a false discovery rate p-value ≤ 0.05, and a fold change ≥|1.5|. A heat map of differentially expressed transcripts was calculated using log counts per million that have been mean-centered and scaled to unit variance. Functional enrichment analysis was performed using Ingenuity Pathway Analysis (IPA) (QIAGEN Inc., Hilden, Germany) to determine the biological functions of significant differentially expressed transcripts. The most significant functional pathways (−log10(p-value) ≥ 1.3) were then selected. Following the selection of canonical pathways related to neuroinflammation, the molecules in those pathways were utilized for IPA network analysis.

## Acknowledgements

We gratefully acknowledge Dr. Kimberly Schuenke for her reviews and editing the manuscript. We thank Dr. Shao-Jun Tang for the consulting and advising, Jill K. Thompson for RNA sequencing. We thank the Sealy Center for Structural Biology and Molecular Biophysics at the University of Texas Medical Branch at Galveston for providing research resources. This work was supported by Pilot grant from University of Texas Medical Branch, Center for Interdisciplinary Research in Women’s Health (CIRWH) and Sealy Center on Aging (SCOA) and the Claude D. Pepper Older Americans Independence Center P30 AG024832 (S.Y.), NINDS R01NS122571 (S.Y.), and R21AI154211 (B.G.). This work was supported in part by the Division of Intramural Research, NIAID, NIH to T. S.

## Author’s contribution

J. L. performed the mouse behavior tests. J. B. performed EV experiments. A. J., J. S., I. G., and C. H. contributed to animal experiments. Y. Q. performed the AFM experiments. Q. C. and T. B. S., contributed to the EV experiments. H.H. performed the RNA sequencing. K.A. and V. T. analyzed sequence data. K. K. conceptualized the bioinformatic data analysis. B. G. conceptualized and designed the EV experiments and prepared the manuscript. S. Y. conceptualized and designed all the experiments, invented the dry ice vapor cold test and optimized test condition, and prepared the manuscript.

